## Supplemental Figure S1-7 for "Discriminative feature of cells characterizes cell populations of interest by a small subset of genes"

**Figure S1_Fujii**

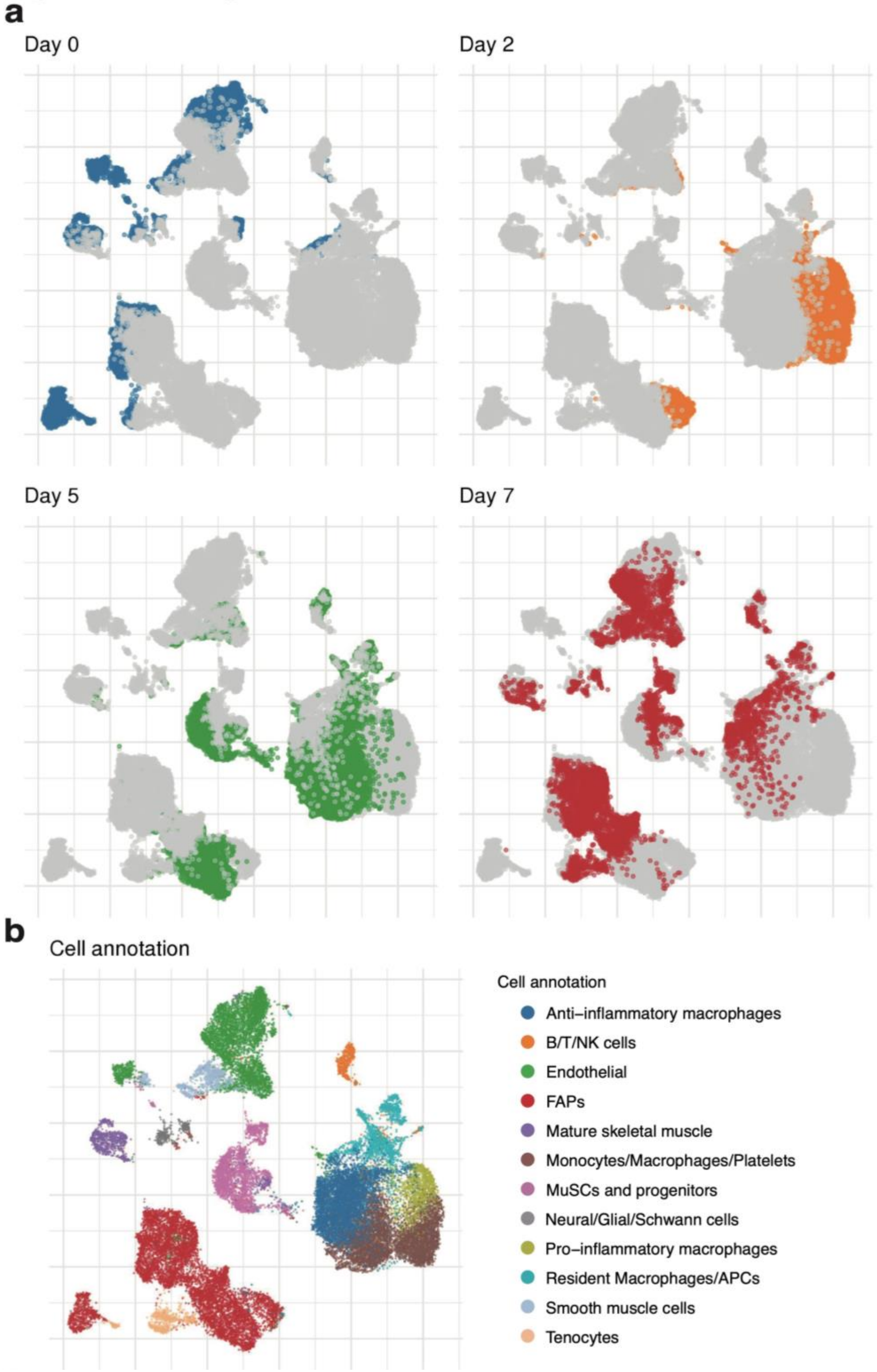


Fig. S1: Original annotations of scRNA-seq data by the authors [GSE143437]. (a) Days after muscle injury. (b) Cell type annotations.

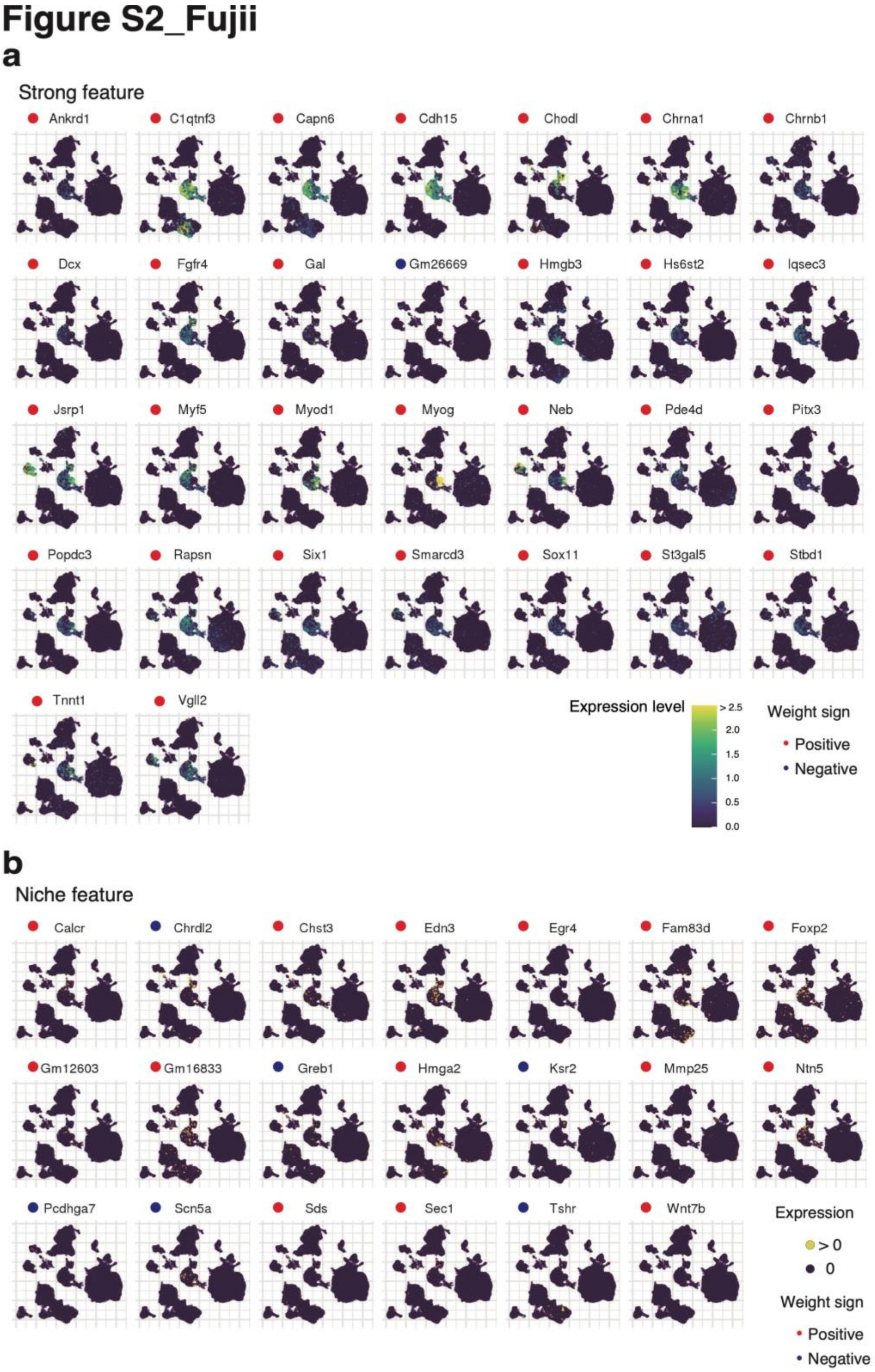


Fig. S2: UMAP visualizations of (a) Strong and (b) Niche features. The markers colored in red/blue indicate the sign of the weight (coefficient) estimated by adaptive LASSO.

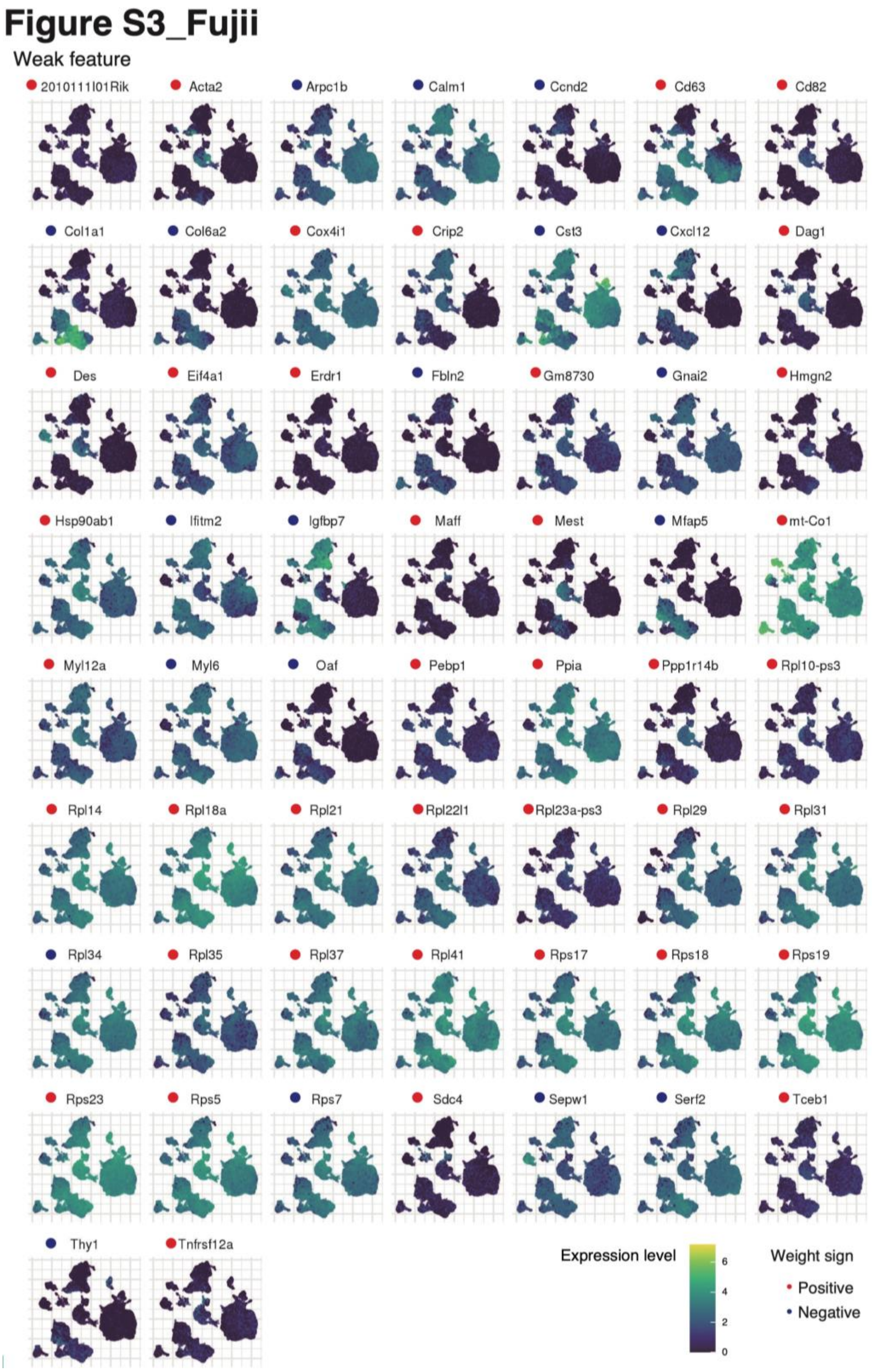


Fig. S3: UMAP visualizations of Weak feature. The markers colored in red/blue indicates the sign of the weight (coefficient) estimated by adaptive LASSO.

**Figure S4_Fujii**

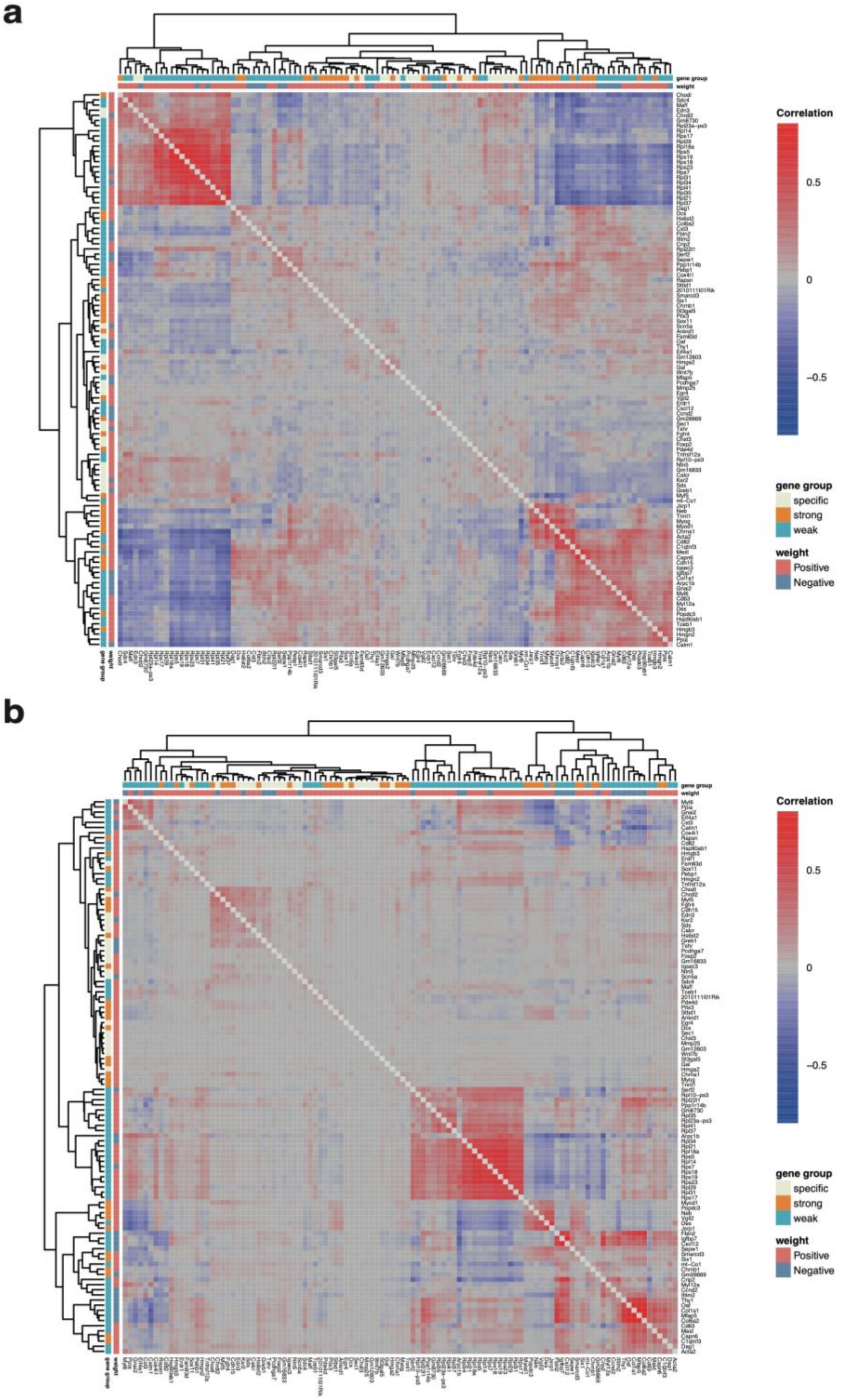


Fig. S4: The correlation coefficient matrix of genes in DFC shows a distinct hierarchical structure within the POI. The matrix in (a) POIs and in (b) Others were shown. Type of feature (either of Strong, Weak or Niche) and the signs of LASSO weight were also indicated.

### Figure S5_Fujii

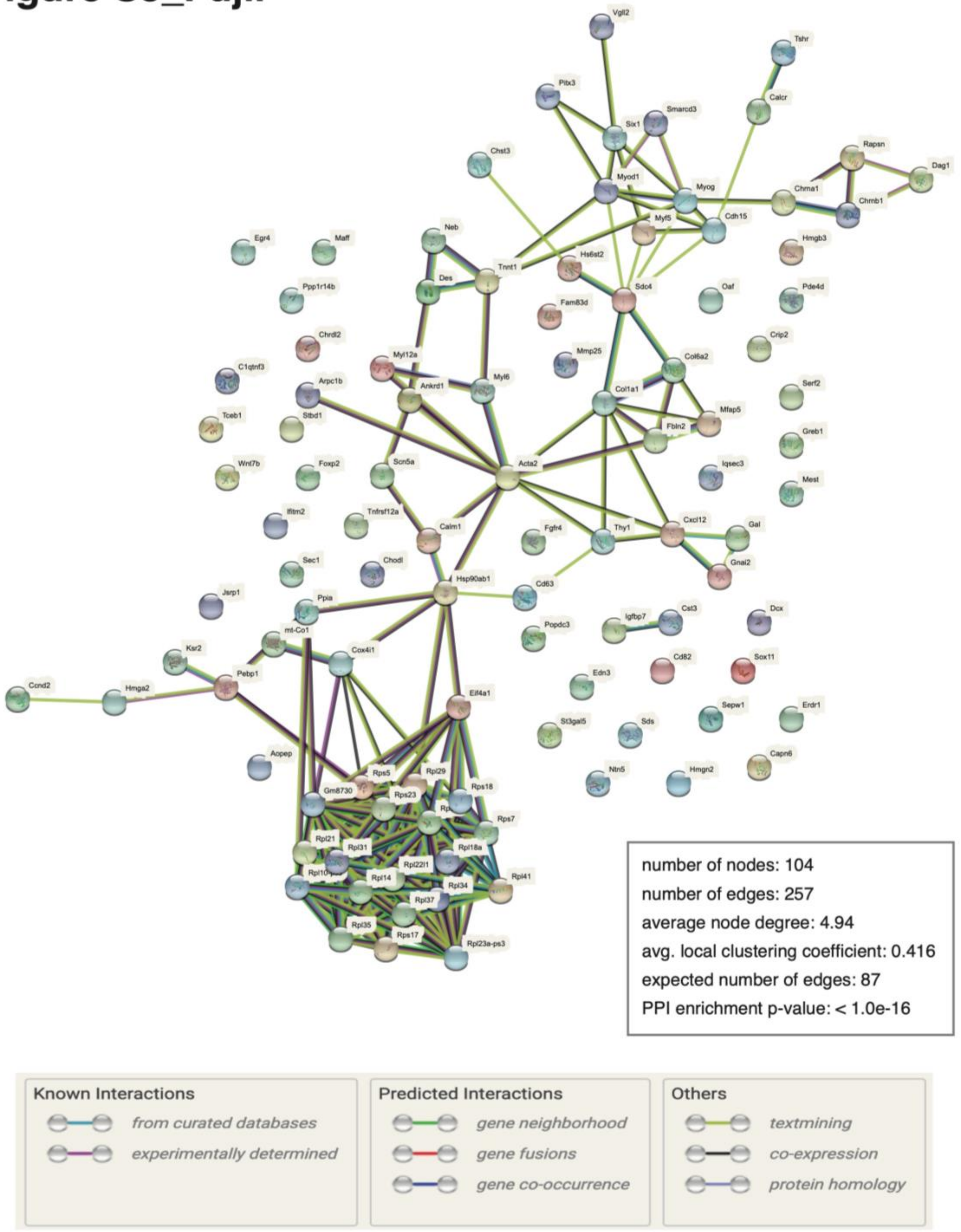


Fig. S5: Functional association of genes in DFC. The functional associations in DFC 108 genes were annotated using STRING.

### Figure S6_Fujii

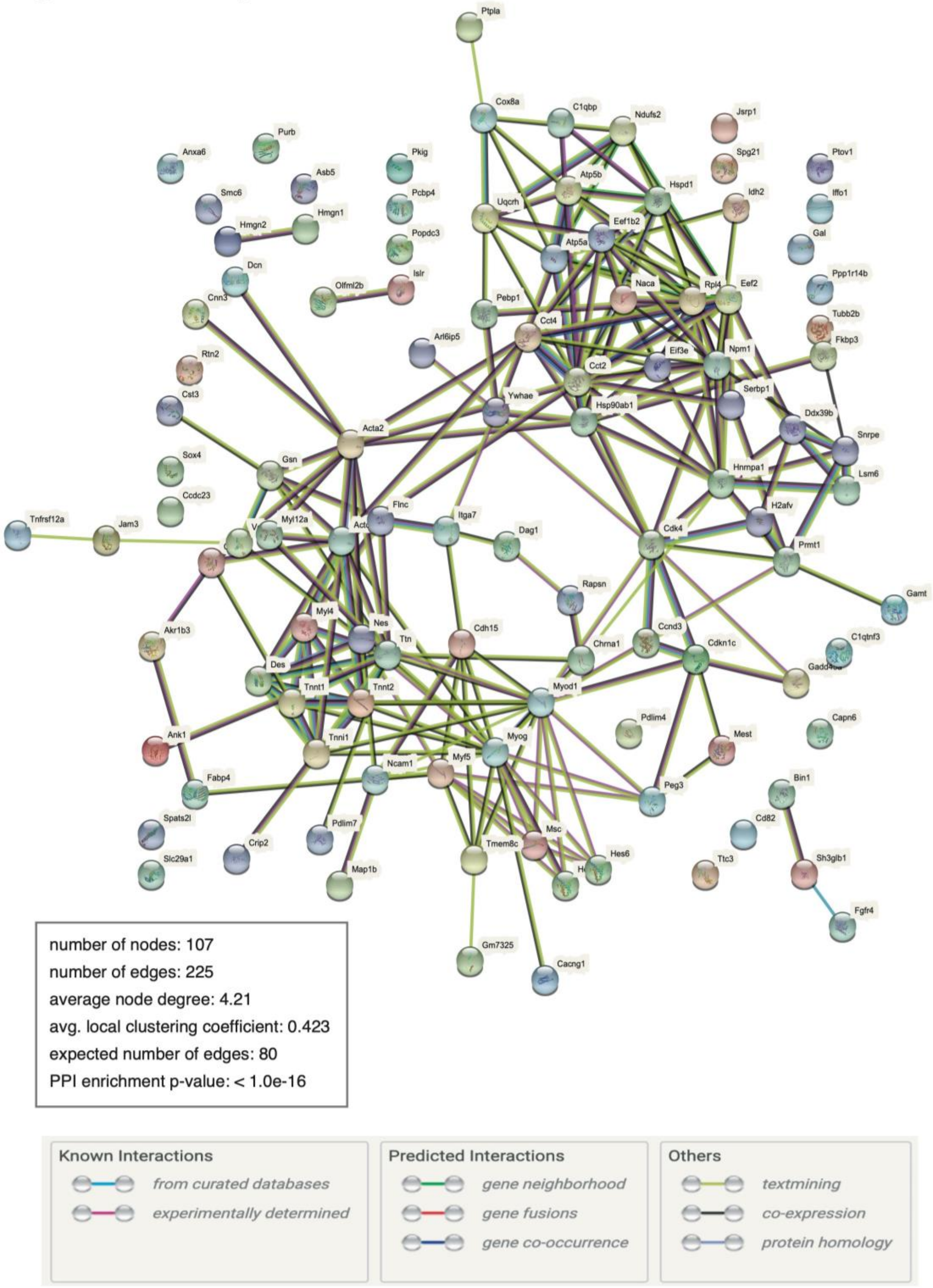


Fig. S6: Functional association of DEGs. The functional association of top 108 genes of P-value in the DEGs annotated by STRING.

Figure S7_Fujii

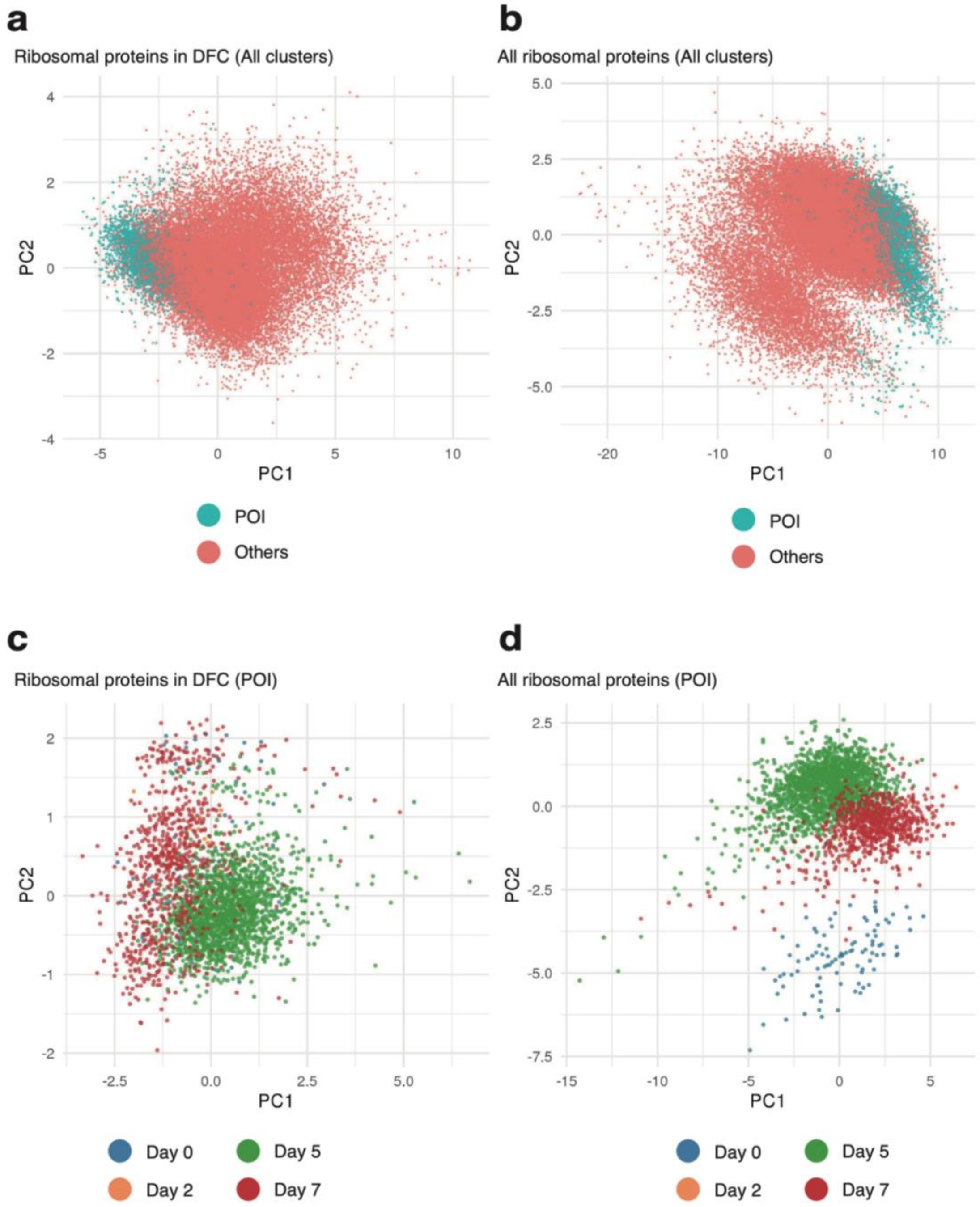


Fig. S7: A certain composition of the ribosomal protein coding genes characterized POI. Scatter plot showing the results of PCA performed using (a) ribosomal protein coding genes in DFC and (b) all ribosomal protein coding genes. (c) PCA performed in POI using genes in (a), and in all cells (POI and others) using genes in (b).
